# Accelerating crystal structure determination with iterative AlphaFold prediction

**DOI:** 10.1101/2022.11.18.517112

**Authors:** Thomas C. Terwilliger, Pavel V. Afonine, Dorothee Liebschner, Tristan I. Croll, Airlie J. McCoy, Robert D. Oeffner, Christopher J. Williams, Billy K. Poon, Jane S. Richardson, Randy J. Read, Paul D. Adams

## Abstract

Experimental structure determination can be accelerated with AI-based structure prediction methods such as AlphaFold. Here we present an automatic procedure requiring only sequence information and crystallographic data that uses AlphaFold predictions to produce an electron density map and a structural model. Iterating through cycles of structure prediction is a key element of our procedure: a predicted model rebuilt in one cycle is used as a template for prediction in the next cycle. We applied this procedure to X-ray data for 215 structures released by the Protein Data Bank in a recent 6-month period. In 87% of cases our procedure yielded a model with at least 50% of C_α_ atoms matching those in the deposited models within 2Å. Predictions from our iterative template-guided prediction procedure were more accurate than those obtained without templates. We suggest a general strategy for macromolecular structure determination that includes AI-based prediction both as a starting point and as a method of model optimization.

## Introduction

The development of AI-based methods for prediction of protein structures has been widely recognized as a turning point for structural biology^1-4^. Predictions using AlphaFold^2^, RoseTTAFold^1^, and related methods^5^ are far more accurate than previous generations of predictions^6^, enabling large-scale analyses of protein function without requiring experimental structural information for each protein^4,7^. Nevertheless, there is considerable discussion about limitations of AI-based models^8,9^.

The potential for using AlphaFold predictions to facilitate structure determination by X-ray crystallography and cryo-EM has been rapidly appreciated in the structural biology community^10-13^. AlphaFold predictions made in the CASP-14 blind test of structure prediction were shown to be effective as starting models for X-ray structure determination using the method of molecular replacement^10^. AlphaFold models and the accompanying PAE (predicted aligned error) matrices can be used to identify domain boundaries in proteins^14^ that could be helpful in designing trimmed versions of proteins suitable for crystallization^15,16^.

AlphaFold prediction can include information from a template, and we recently showed that a prediction based on a template can be more accurate than either a sequence-based prediction or the template itself^17^. This property allows iterative improvement of modeling and AlphaFold prediction, and we here describe an automated procedure that can accomplish this using X-ray crystallographic data.

## Results

### Crystallographic structure determination using iterative AlphaFold prediction and rebuilding

Our iterative procedure for macromolecular structure determination by X-ray crystallography uses AlphaFold predictions in an initial structure solution cycle, followed by cycles of AlphaFold prediction and model rebuilding in which rebuilt models from one cycle are used as templates for prediction in the next^17-19^ (see Methods). This procedure requires processed crystallographic data (structure factor amplitudes and their uncertainties, space group and unit cell dimensions for the crystal, and the resolution of the data) and information about the sequences of the macromolecules present in the crystal. It produces an optimized (density-modified^20^) electron density map and a model based on that map. During the procedure, the map itself improves as the model improves, in contrast to the related workflow for single-particle cryo-electron microscopy^17^, where the map does not change. The approach can be used if the majority or all of a structure consists of protein. If other components are present, such as RNA/DNA, only the protein part of the resulting map is interpreted.

### Determination of challenging structures using AlphaFold prediction

The method of molecular replacement^21^, in which an initial model that is similar to the structure to be determined is used as a hypothesis for the actual structure, has been applied in about 80% of recent macromolecular crystal structure determinations^22^. Once the orientation and location of the initial model is found, the model and density map are usually improved by cycles of map calculation alternating with map representation as an atomic model with restrained geometry^23^. These procedures work best if the initial models are within about 1 Å to 2 Å (rmsd of C_α_ atoms) over about 50% of the structure^24^. Excitement about AlphaFold predictions in the crystallographic field comes from the observation that such predictions may generally be accurate enough to provide the necessary starting point for molecular replacement, largely removing the need for using anomalous scattering or other experiments to obtain crystallographic phases^10,13,25^.

Here we use an automated procedure to test the iterative use of AlphaFold predictions in macromolecular crystallography. To recreate a situation where challenging new structures are being determined with AlphaFold predictions, we selected structures obtained with anomalous scattering, an approach typically used when molecular replacement is expected to fail, but we did not include the anomalous diffraction information. For Protein Data Bank^26^ (PDB) entries released in the 6-month period starting from Dec. 8 2021 through June 29, 2022, this selection yielded 215 unique structures with resolutions ranging from 1.0 Å to 4.6 Å.

We applied our iterative procedure for AlphaFold prediction and model improvement to each of the 215 deposited datasets. In seven cases, the initial molecular replacement step did not yield any solution and the analysis was not continued. For the remaining 208 datasets our procedure generated density-modified^20^ electron density maps and models interpreting those maps. To identify which maps and models were at least partially correct, we compared the density-modified electron density maps with model-based (*F*_*calc*_) maps calculated from the corresponding deposited structures. Using a conservative minimum map correlation of at 0.5 as a threshold^27^, 187 of 215 analyses (87%) were successful and the remaining 28 (including the seven that failed in molecular replacement) were unsuccessful.

Figure 1A shows the distribution of map-model correlation values. For the successful cases, the average map correlation was 0.84. The solid bars in Fig. 1B illustrate the completeness of these structures, as measured by the percentage of C_α_ atoms in deposited models matching those within 2 Å in the rebuilt models^17^. The average completeness was 90%, and models for all but one were at least 50% complete. For the unsuccessful analyses (open bars), the completeness of the models was much lower (average of 20%). In a few of the successful instances the final structures contained domains that matched the deposited model, but the connectivity between domains was incorrect (e.g., PDB entry 7E1D which is a domain-swapped^28^ dimer). We included all matching parts by using space-group symmetry-related copies of each chain from the deposited models in the comparisons. The overall high success rate shows that in most cases, AlphaFold predictions are accurate and complete enough to be used as starting models in macromolecular crystallography.

**Fig. 1.**
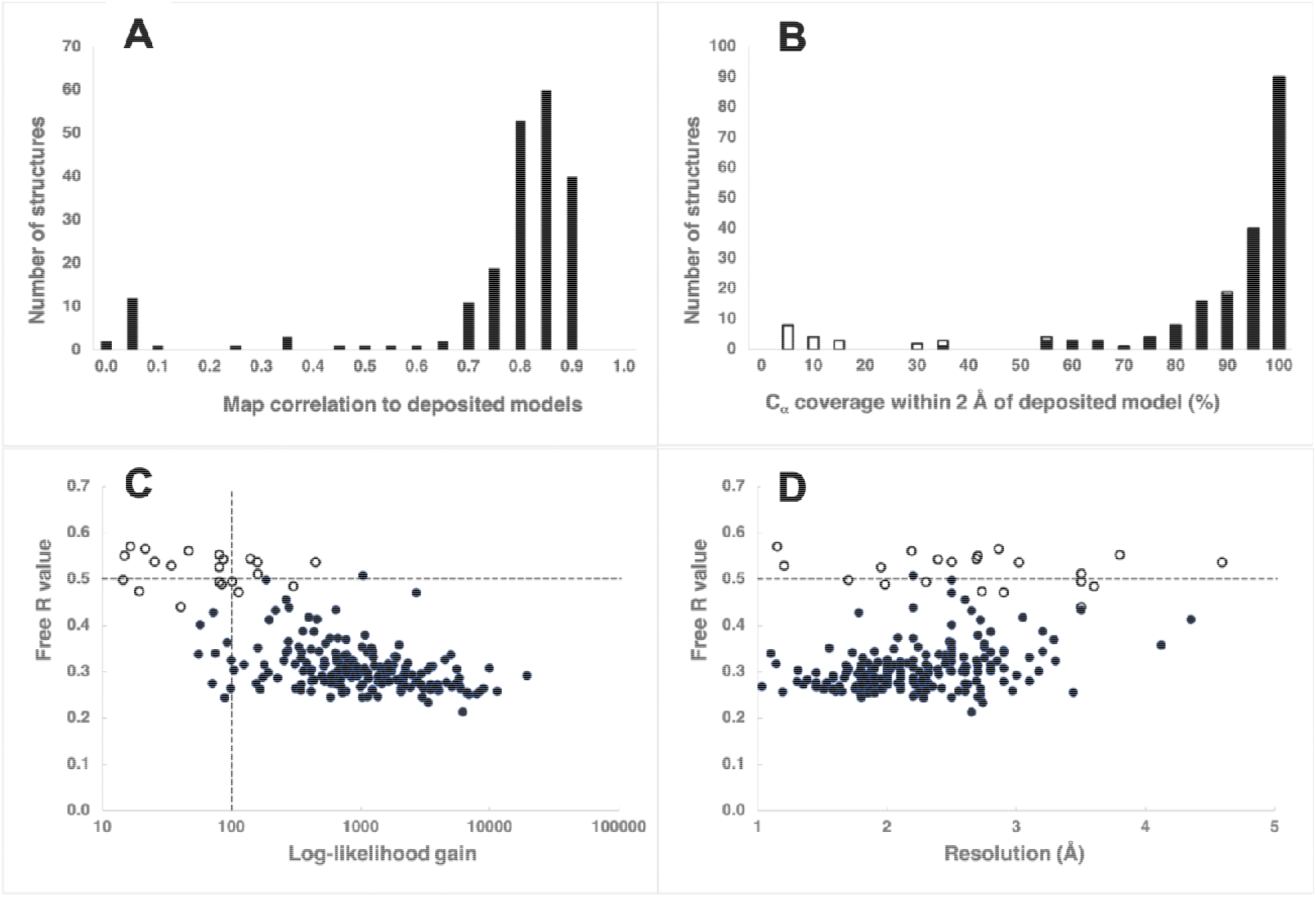
Structure redeterminations using AlphaFold predictions obtained during automated structure redeterminations for the 208 analyses that yielded a molecular replacement solution. A. Correlation of maps calculated from deposited models to density-modified maps^20^. B. Percentage of C_α_ atoms in redetermined models within 2 Å of a C_α_ atom in deposited models, after applying overall space-group specific origin shifts and including space-group symmetry-related copies in the comparison. Successful cases are shown with solid bars and unsuccessful cases are shown in open bars. C. Log-likelihood gain in molecular replacement and free R values for automatic structure analyses. Unsuccessful cases are indicated by open circles. D. Free R values as in C, except shown as a function of the high-resolution limit of the X-ray data.

AlphaFold provides residue-level confidence estimates including a predicted Local Difference Distance Test (pLDDT) associated with each amino acid residue^2^. Here we refer to residues with pLDDT values of 90 or higher as “high confidence”, and those with values of 70 or higher as moderate-to-high confidence^2^. The success of automated structure determination using AlphaFold prediction was strongly dependent on the percentage of moderate-to-high confidence residues. For successful cases, an average of 90% of residues in the AlphaFold predictions were predicted with moderate-to-high confidence, and the smallest percentage of moderate-to-high confidence residues was 30%. For the 28 unsuccessful cases, the average percentage predicted with moderate-to-high confidence was much lower (50%).

We assessed the similarity of predicted models and corresponding deposited structures in more detail by superimposing the models, removing residues with low confidence (pLDDT < 70), and further removing residues where the rmsd (smoothed with a window of 10 residues) was greater than 3 Å. We then noted the rmsd between matching C_α_ atoms and the coverage, defined here as the percentage of residues in the deposited model that were matched by the predicted model. For successful cases, the average C_α_ rmsd was 1.2 Å and the average coverage was 89%. Such values are normally associated with success in molecular replacement^24^. For unsuccessful cases the average rmsd was 2.8 Å, much too high to expect success in molecular replacement, and the average coverage was 50%, a borderline value for success. Overall, 19 of the 28 failures had either a coverage less than 50%, an rmsd over 2 Å, or both.

When determining a new structure, metrics that do not depend on knowing the true structure are important for evaluating whether structure determination has been successful. Fig. 1C shows two metrics that are available after structure determination and that are independent from knowledge of the true structure^19^. Log-likelihood gain (LLG) scores reflect the confidence of placing the model, and free R values reflect the cross-validated agreement of the model with crystallographic data after model rebuilding and refinement. The median LLG score for successful structure determinations with AlphaFold predictions was about 1000, while for unsuccessful cases the median was 80. We note that in a large-scale test of molecular replacement with homology models from the PDB^29^, only 7% of cases had an LLG score over 1000 (median LLG score of 270). Our procedure using AlphaFold predictions therefore led to much higher LLG scores than typically obtained with homology models. The median free R value for successful cases was 0.30, a value normally associated with a largely-correct but unfinished structure (structures that do not contain waters, ligands, or any components other than protein). Fig. 1C illustrates that most unsuccessful solutions had free R values above 0.5, log-likelihood gain scores below 100, or both, suggesting that the free R value and log-likelihood gain can be used effectively to evaluate potential solutions in new structure determinations using this procedure, as is standard practice^19^. Fig. 1D displays the free R values as a function of the high-resolution limit of the X-ray data and illustrates that the range of free R values depends slightly on the resolution of the data.

We evaluated whether our procedure may have yielded partially-correct solutions for some of the cases that we considered unsuccessful. For example, the model obtained by automated structure determination for PDB entry 7f05 had a map-model correlation of just 0.24 and was thus considered not successful. Supplementary Fig. 1A shows that much of the rebuilt chain B for this structure (dark brown) very closely matches chain A in the deposited model (symmetry copy from the deposited model shown in light blue). However, the rest of the rebuilt model does not match the deposited structure. Supplementary Fig. 1B shows that the density-modified map is also quite clear. This example shows that even an unsuccessful structure can contain some correct information. Furthermore, although our procedure did not complete the structure, other approaches such as repeating molecular replacement using this one chain as a starting point^19^ might well do so.

### Iteration of AlphaFold prediction and model optimization

Our procedure for automatic structure determination iterates through AlphaFold prediction, using rebuilt chains obtained at the end of one cycle of iteration as templates for AlphaFold prediction in the next. In this procedure, map and model quality can improve at two stages. First, the new AlphaFold prediction using the previously rebuilt model as a template can yield an improved predicted model^17^, and second, rebuilding can improve both model and map^23^. Iteration is carried out until the change in the model from one cycle to the next is small. As models are not necessarily improved with rebuilding, they are evaluated based on their free R value, i.e. they are kept if the free R value decreases.

Fig. 2 shows this procedure applied to PDB entry 7oa7^30^ which includes X-ray data to a resolution of 1.45 Å. This figure follows the models and their fit-to-map through two cycles of the procedure. The lefthand column shows ribbon overviews of superpositions onto the deposited 7oa7 deposited model (always in cyan). The two righthand columns are closeups superimposed on the map at the relevant stage: the central column (B, E, and H) shows the progression for a region that started out matching the deposited structure quite well, and the righthand column (C, F, and I) shows the progression for a region that started out matching quite poorly. For these two righthand columns, predicted models are in magenta and rebuilt models in dark blue, and time moves vertically downward. The top row shows the initial no-template prediction (magenta) and the map output from molecular replacement. The central row shows the rebuilt (dark blue) model and the map as rebuilt in cycle 1. The bottom row shows the cycle 2 predicted model (magenta) that was given the rebuilt cycle 1 model as a template, and the cycle 2 map.

**Figure 2.**
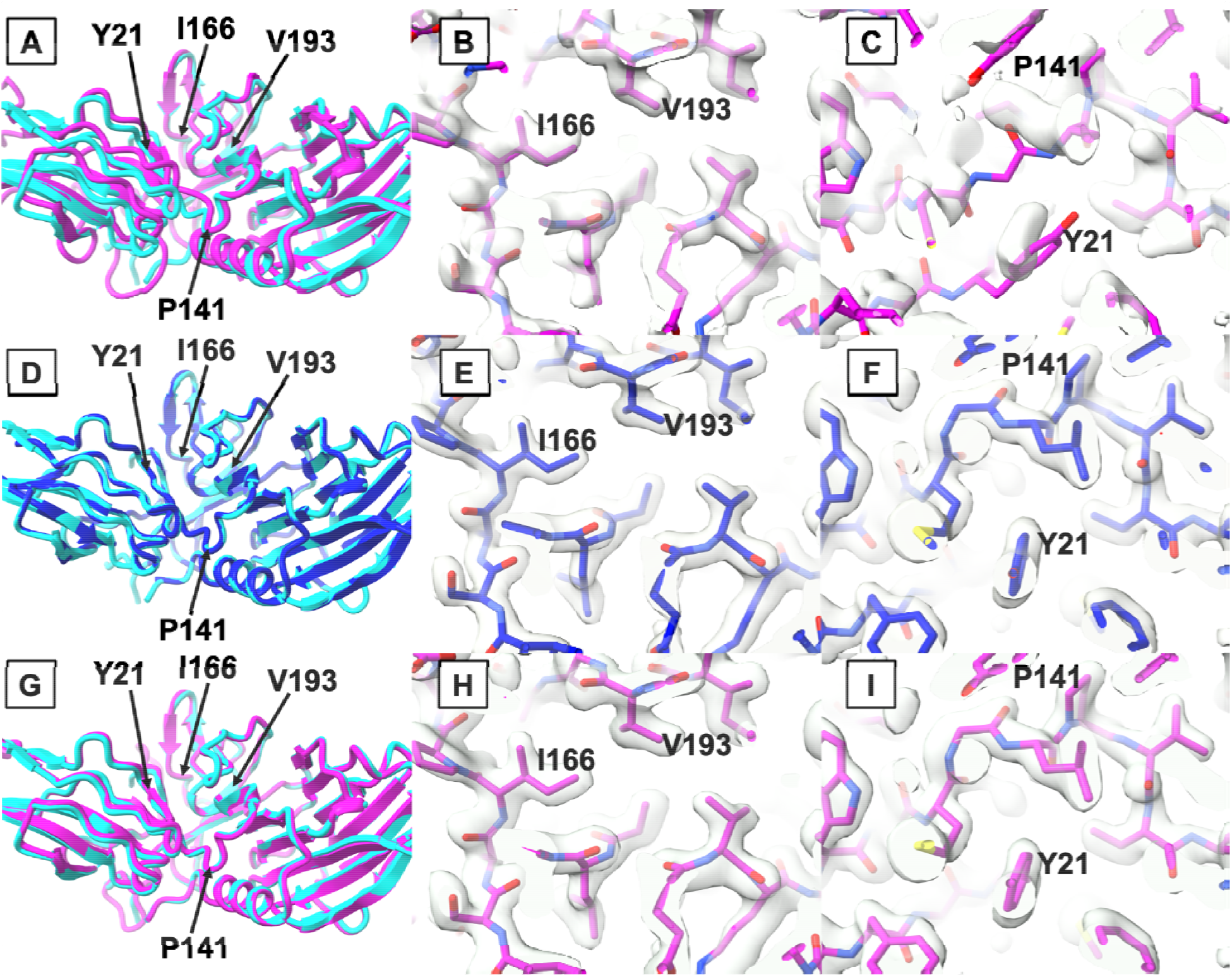
Iterative AlphaFold prediction and model-rebuilding with X-ray data for PDB entry 7oa7. A: Superposition of AlphaFold prediction without templates (magenta) and the deposited model (cyan). The region comprising residues I166 and V193 and the domain on the right are similar in the deposited and predicted models, while the region of residues Y21 and P141 and the domain shown on the left differ between deposited and predicted models. B and C: Close up views of the AlphaFold prediction without templates superimposed on the density map obtained after molecular replacement and refinement. The good region containing residues I166 and V193 is shown in B and the poor region containing Y21 and P141 is shown in C. D superposition of the deposited model and the model obtained by molecular replacement and automated model rebuilding using the AlphaFold prediction shown in A (dark blue). E and F: Close up of the model in D and density-modified map obtained from the first cycle of automated model rebuilding. G: Superposition of the deposited model and superposed AlphaFold prediction obtained using the model in B as a template. H and I: Close up of the predicted model in G and density-modified map obtained in the second iteration of model-rebuilding.

An initial AlphaFold prediction obtained using the sequence but no structural templates is shown in Fig. 2A. Some parts of the predicted model (magenta) match the deposited model (cyan) quite closely, such as the domain on the right side of Fig. 2A and the region near residues I166 and V193. Other parts, such as the region near residues Y21 and P141, have different backbone conformations. Still other parts, such as much of the domain on the left side of Fig. 2A, differ by a combination of local conformations and overall rotation and translation. Overall, the predicted model differs from the deposited model by a C_α_ rmsd of 2.9 Å. The initial density map obtained after molecular replacement is quite clear in the region of residues I166 and V193 where the predicted model matches the deposited model closely (Fig. 2B). In contrast, the map is less clear and does not match the predicted model in the region of residues Y21 and P141 where the deposited and predicted models differ (Fig. 2C).

After automated crystallographic rebuilding starting with the predicted model shown in Fig. 2A, the rebuilt model (dark blue) improves substantially (Fig. 2D); the overall C_α_ rmsd to the deposited model is 0.1 Å, and 90% of C_α_ atoms in the deposited model are within 2 Å of those in the rebuilt model. The density map is also considerably improved (Fig. 2E and 2F) and matches the rebuilt model quite closely in the region near I166 and V193 and, most notably, in the region near Y21 and P141 where the predicted and deposited models differed in the previous step.

Using the rebuilt model (Fig. 2D) as a template, a new AlphaFold prediction was obtained that is very similar to the deposited model (Fig. 2G, overall C_α_ rmsd of 0.5 Å) and that is quite different from the initial AlphaFold prediction (Fig. 2A). Figs 2H and 2I show that the template-based prediction matches the density-modified map, both in the region near I166 and V193 where the initial predicted model matched the deposited model and in the region near Y21 and P141 where it did not. The template-based prediction is even more complete than the template, with 99% of residues in the deposited model matched within 2 Å by a C_α_ atom in the prediction. (The model used as a template had only 90% matched residues). Note that both the polypeptide backbone and the side chains in the template-based prediction closely match the density map (Fig. 2I) and corresponding side-chain positions in the rebuilt model (compare Fig. 2F and Fig. 2I)

Two additional cycles of rebuilding based on the model in Fig. 2C resulted in a 95% complete model, i.e., where 95% of C_α_ atoms in the deposited model matched those within 2 Å in the rebuilt model, with an overall C_α_ rmsd of 0.1 Å.

In summary, though the initial AlphaFold prediction obtained for PDB entry 7oa7 was substantially different from the deposited model, a structure could be obtained on the first cycle of our procedure with standard molecular replacement and rebuilding procedures. Using the rebuilt model as a template for AlphaFold prediction, a new prediction was obtained that was far more accurate than the initial prediction. Three cycles of iteration for PDB entry 7oa7 led to a nearly-complete model matching the deposited structure closely, and required approximately 7 hours using 4 processors on a Linux server.

Although the template-based AlphaFold prediction for PDB entry 7oa7 is more complete than the rebuilt model, it is slightly less accurate (C_α_ rmsd with deposited model of 0.5 Å vs 0.1 Å for Alphafold and rebuilt model, respectively). This is not surprising, as the AlphaFold prediction uses the experimental data only indirectly (in the form of the rebuilt model) and has not been adjusted to match the density map. In a real case, both the rebuilt model and final AlphaFold prediction would have utility in subsequent stages of structure determination, as the rebuilt model is slightly more accurate and the AlphaFold model is more complete.

Fig. 3A shows histograms of free R values on the first and last cycles of our procedure for all 187 successful analyses. The free R value decreased with iteration in 88 of 187 cases, and on average, it was reduced from 0.33 to 0.31. Iteration was more successful when the free R value after the first cycle was poor. For the 109 cases where the free R value on the first cycle is worse than 0.30, 66 were improved by iteration, while for the 78 cases where the free R value was better or equal to 0.30, only 22 were improved. Presumably this is because the procedures used here do not generally yield free R values much better than 0.30.

**Figure 3.**
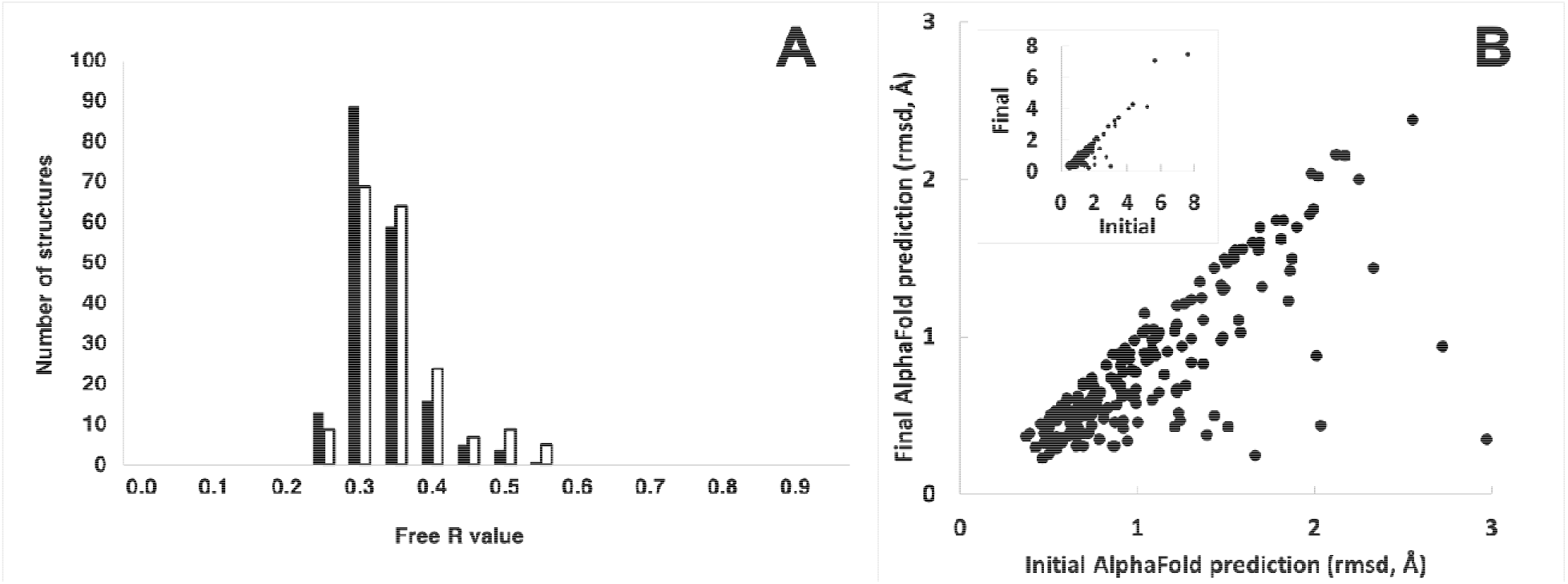
Iterative AlphaFold prediction and model-rebuilding with X-ray data. A: Free R values for the rebuilt model after the first (open bars) or last (closed bars) iteration of prediction and rebuilding (using the cycle with the lowest free R value as the last iteration). B: Rms differences between the first chain in each deposited model and corresponding initial AlphaFold predictions (carried out without templates at the beginning of the first cycle, abscissa), and final AlphaFold predictions (carried out with templates on the last cycle, ordinate). One PDB entry (7dz9) is not shown as no rebuilt model was obtained for the first chain in the PDB entry so that the AlphaFold models on each cycle were the same. Inset includes 9 cases not shown in the main panel where the initial rmsd was greater than 3 Å.

Fig. 3B shows the rmsd between C_α_ atom coordinates in AlphaFold predictions for each structure and those in the corresponding chains of deposited models. Most points are below the diagonal indicating that the final AlphaFold prediction was more accurate than the initial one. For initial AlphaFold predictions, the median rmsd was 1.0 Å, while for final predictions with templates it was reduced to 0.7 Å.

Overall, Fig. 3 shows that the agreement of rebuilt models with experimental data was generally very good at the end of the first cycle (mean free R of 0.33), and improved slightly with iteration (mean free R of 0.31). The accuracy of initial AlphaFold predictions was also good, and improved substantially with iteration using rebuilt models as templates.

### Characteristics of rebuilt and predicted models

We examined the geometric characteristics of four sets of models: initial AlphaFold predictions without templates, final-cycle AlphaFold predictions that used a template, the final rebuilt model and the deposited model. Fig. 4A, 4B and 4C show comparisons between the initial AlphaFold prediction without template and the final rebuilt model, and Fig 4D, 4E, 4F show comparisons between the AlphaFold predictions with templates and the deposited model. Fig. 4A shows the distribution of percentages of residues in favored Ramachandran conformations^31^. The AlphaFold predictions (open bars) have a mean percentage of 98% while the rebuilt models (solid bars) average to 96%. The percentages of rotamer outliers^31^ (Fig 4B) are 0.3% and 2 % for Alphafold (open bars) and rebuilt models (solid bars), respectively. Clashes between non-bonded atoms were somewhat worse in the AlphaFold predictions (Fig. 4C, open bars, mean Clashscore^32^ of 29) than for rebuilt models (solid bars, mean Clashscore of 10). Fig. 4D shows the percentage of favored Ramachandran conformations for deposited models (closed bars, mean of 97%) and AlphaFold predictions using a template (open bars, mean of 98%). Fig 4E shows rotamer outliers for deposited (closed bars, mean of 2%) and AlphaFold predictions with templates (open bars, 0.3%), and Fig. 4F shows Clashscore values for deposited models (closed bars, mean of 6) and AlphaFold predictions with templates (open bars, mean of 27). Overall, the AlphaFold predictions (with or without templates) have somewhat more favorable main-chain Ramachandran conformations and the fewest rotamer outliers of the four sets of models, while the deposited models have the best Clashscore values.

**Fig. 4.**
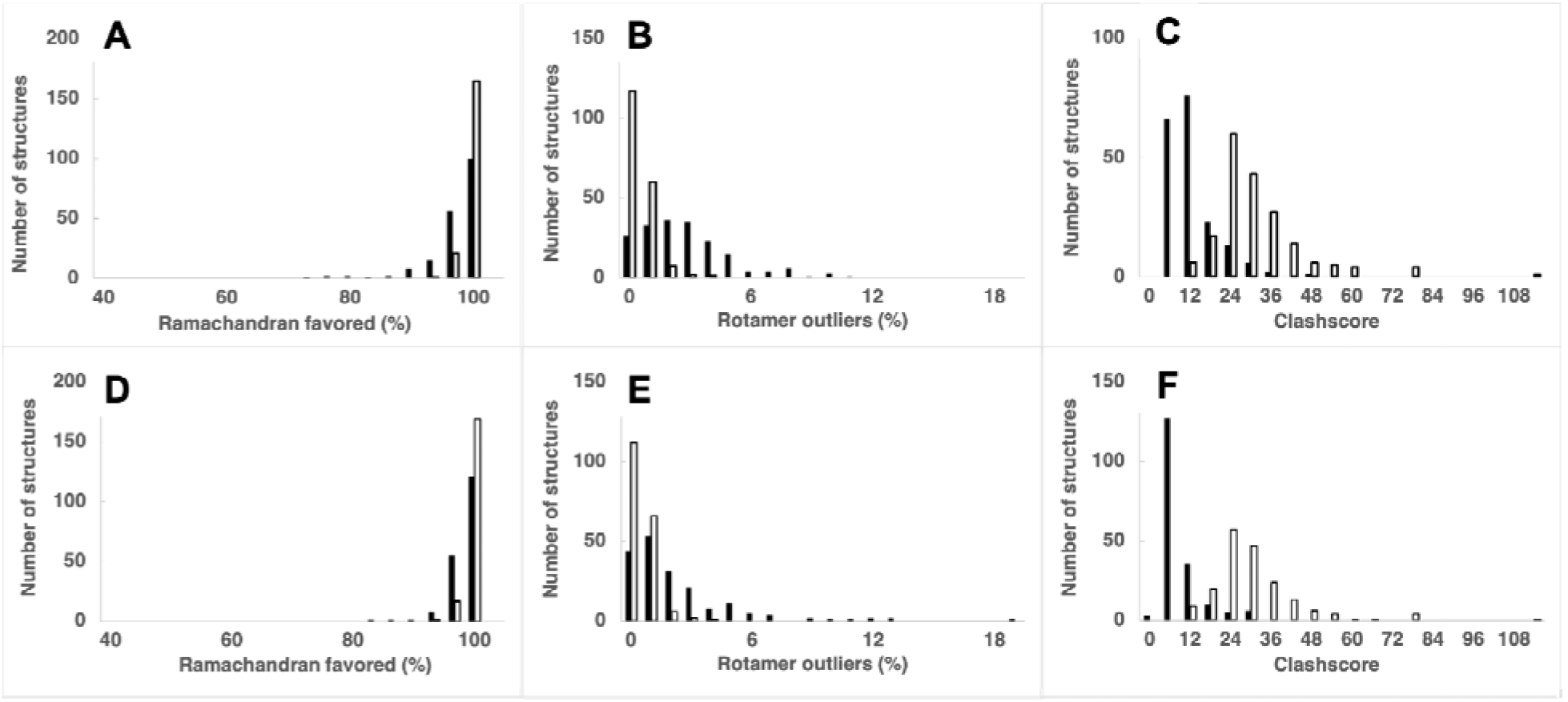
Geometric and packing characteristics of predicted and rebuilt models. A: Histogram of the percentage of residues with “favored” backbone conformations (open bars, AlphaFold predictions obtained without templates, closed bars, rebuilt models). Calculations for AlphaFold predictions include residues with pLDDT of 70 or above. B: Histograms of rotamer outliers (unlikely side-chain conformations). C: Histograms of clashscore^32^ values. D, E, and F, as A, B, and C except that closed bars represent deposited models and open bars represent AlphaFold models with templates (final cycle of iteration, trimmed to remove residues not matching map or with pLDDT of less than 70, see Methods).

## Discussion

### Strategy for crystal structure determination using template-guided AlphaFold prediction

Our analysis shows that AlphaFold predictions obtained based on sequence information alone are usually accurate enough to solve the crystallographic phase problem with molecular replacement. As we have shown previously, partially-complete rebuilt models, such as those obtained by automatic model-building, can be used effectively as templates to guide subsequent AlphaFold predictions, and the resulting predictions can be more accurate than either the template or predictions made without templates^17^. Finally, we find that if a relatively accurate model is used as a template for AlphaFold prediction, the resulting predicted model can maintain both the overall conformation of the template and details of side-chain conformations.

These observations suggest how the overall strategy for macromolecular crystal structure determination can be revolutionized with AI-based predictions. In this new strategy, AlphaFold predictions are considered to be hypotheses for the actual structure. These predictions have confidence measures reflecting the expected accuracy for each residue in the prediction. Initially, the information available for structure prediction consists of the sequence for each chain and multiple sequence alignments based on those sequences. As structural information is accumulated, new predictions are made that incorporate this information.

This new strategy for macromolecular crystal structure determination starts at the planning stage and continues through the stage of obtaining a final model. In the planning stages of an experiment, confidence measures for AlphaFold predictions can be used to assess whether predicted models obtained using sequence alone will be useful as starting points for structure determination. Also, the AlphaFold predictions can be used to design trimmed versions of a macromolecule that can be successfully crystallized^16^.

To begin structure solution, AlphaFold predictions, trimmed to remove low-confidence regions, can be used as search models in molecular replacement, with a high probability of yielding a molecular replacement solution that is at least partially correct.

As in a conventional structure determination workflow, the next stage consists of refinement of the molecular replacement solution, followed by iterations of map calculation, density modification, and model rebuilding. This stage is typically carried out by existing automatic procedures that can improve both the model and the accuracy of the density maps^23^. If there are major components of the structure that are not protein (i.e., RNA or DNA), these components would need to be built based on the density map, yielding a more complete model and an improved density map as well. In our procedure, such additional components are automatically used in subsequent cycles.

Once an improved model is obtained, AI-based prediction can be used again. This time, a working model is used as a template to guide AlphaFold prediction, as demonstrated here. This step can be thought of as a method of model optimization using AlphaFold. The working model guides AlphaFold prediction, yielding a prediction that has information from the working model but potentially with improved geometry or a more accurate overall conformation. This template-guided AlphaFold prediction can be carried out with or without information from multiple sequence alignments. The optimized models obtained in this way can be quite accurate, but as they are obtained without direct use of experimental information, they may not exactly match the density map at this point.

The new set of AlphaFold predictions can then be superimposed on corresponding parts of the working model, refined using experimental data, and then considered as new hypotheses about the structure. It is anticipated that after a typical application of the procedure described here, both the working model and the refined new AlphaFold predictions would be compared with the density map, and a new composite model incorporating the best parts of each would be created. This composite model could be used as the starting point for another iteration of model rebuilding, or if it is sufficiently complete, it could be used as a starting point for adding ligands, waters and other small molecules, and covalent modifications. If the resolution of the data is high, this model might also be modified to explicitly add alternate conformations.

This overall strategy for structure determination differs from long-established procedures at three important steps: (1) at the planning stage where AlphaFold predictions can give an idea of how challenging an experiment will be and where they can guide the design of crystallization constructs^16^, (2) at the initial structure determination stage where trimmed AlphaFold predictions can be used in molecular replacement^10,11,13^, and (3) at the model optimization stage where template-guided AlphaFold prediction may be able to improve models continuously in the analysis^17^.

A central part of this overall strategy is making use of the synergy between model rebuilding and AlphaFold prediction. Using a model obtained in one cycle as a template for AlphaFold in the next cycle often leads to improvement in AlphaFold prediction, which can result in better model-building and density maps in the next cycle as well. Iteration of this procedure therefore can yield improved AlphaFold predictions, final rebuilt models, and density maps. According to our analysis, improvement of AlphaFold prediction is more pronounced (Fig. 3B) than the improvement in density maps (as reflected in the free R value, Fig. 3A). This is likely due to our use of long-established and very effective procedures for iterative model rebuilding with macromolecular crystallographic data during the first cycle, so that subsequent cycles have less effect on the density map and have a larger effect on the AlphaFold predictions.

We have previously taken advantage of iterating AlphaFold prediction and model rebuilding in the interpretation of density maps from cryo electron microscopy (cryo-EM)^17^. The synergy between AlphaFold prediction and model-building is similar in the crystallographic and cryo-EM cases, but not identical. The major difference is that in the case of crystallographic data, the density map can improve dramatically, while with cryo-EM data, the density map is generally fixed. Although it might be anticipated that this would result in better AlphaFold predictions and working models by iteration with crystallographic data, our tests show that the improvement is similar for the two types of experimental methods. For crystallographic data, the median rmsd between AlphaFold predictions and matching PDB entries was reduced from 1.0 Å to 0.7 Å by iteration; for cryo-EM structures, the initial median rmsd was higher (2.5 Å), but was reduced proportionally to 1.6 Å by iteration. It seems possible that in both cases the limiting step may be model rebuilding, and with recent developments in AI-based map interpretation^33^ this limitation might be greatly reduced, potentially leading to more improvement than already obtained by iteration.

Our tests of an automatic procedure for initial crystallographic structure determination demonstrate that this overall strategy is likely to be generally effective, as most of the challenging structures we analyzed could successfully be redetermined.

## Acknowledgments

The authors appreciate funding from Lawrence Berkeley National Laboratory (grant DE-AC02-05CH11231 to PDA), from the Phenix Industrial Consortium (to PDA), from the National Institutes of Health (grant GM063210 to PDA, RJR, TCT, JSR) and from the Wellcome Trust (grant 209407/Z/17/Z to RJR).

## Author contributions

The overall concepts in the paper were developed and supervision was carried out by TCT, PDA, RJR, and JSR. TCT wrote the initial draft and carried out the analyses. AJM, BKP, PVA, TIC, CJW, DL and RDO developed tools that were essential to the work. All authors contributed ideas to the work and assisted in editing of the manuscript.

## Methods

### Data and Models from the PDB

We chose models from the PDB with a goal of obtaining a representative set of challenging structures determined after the training of AlphaFold (which used structures through April, 2018). We selected all 215 unique protein-containing structures determined by the method of single wavelength anomalous diffraction (SAD) over a 6-month period (release dates from Dec. 8, 2021 to June 29, 2022). Crystallographic data for each model were obtained from the PDB. The X-ray data typically included the anomalous data used to solve the structures, but we did not use the anomalous information (Bijvoet pairs of reflections were averaged). Some structures contained non-protein contents, such as waters, ions, small molecules, RNA, DNA, covalent modifications. All of these were removed from deposited models and were not considered in our analyses. For structural comparisons, protein chains from deposited models were superimposed on our automatically-generated models using crystallographic symmetry operations (with origin shifts, as appropriate).

### Automatic structure determination with iterative AlphaFold prediction

Our procedure for automated structure determination consists of cycles of AlphaFold prediction, trimming and splitting predictions into compact domains, molecular replacement (on the first cycle), morphing full-length predictions onto the model obtained from molecular replacement or from a previous cycle, refinement, model rebuilding and trimming. These steps are described in detail below. At least three cycles are carried out, and the process is terminated when the rebuilt models for subsequent cycles have an overall rmsd to the previous model of less than 0.25 times the high-resolution limit of the X-ray data (controlled by the parameter *cycle_rmsd_to_resolution_ratio*). The rationale for scaling this to the resolution is that lower-resolution structures are anticipated to have larger coordinate errors. The entire process is completely automated and can be carried out with the *Phenix* tool *PredictAndBuild*. We used default parameters in all the structure redeterminations described here.

### AlphaFold prediction

We used a local installation of AlphaFold2^1^, configured as a server using software from ColabFold^34^, to carry out AlphaFold predictions using simplified methods available in *Phenix*^*35*^. The sequences used in prediction were obtained from the sequence file supplied by the PDB for the corresponding structure. An alternative to using a local installation is available in *Phenix*; this alternate method uses Google Colab with a *Phenix* script to carry out predictions.

Predictions without templates were carried out using random seeds to initiate five AlphaFold predictions; the prediction with the highest average pLDDT (confidence) value was kept.

Predictions with templates were carried out in the same way, but supplying a template. Templates consisted of the rebuilt model obtained on a previous cycle of prediction and rebuilding. Two forms of templates were used, one containing just main-chain and C_β_ atoms, and one including all side-chain atoms. For shorter chains both forms of the template were used (10 total predictions) and for longer ones only the main-chain and C_β_ atoms were included. This limitation was due to our server and the uploading method for template files; side-chain-containing templates packaged for uploading could not have more than 65536 characters; this limitation is not present when using the *Phenix* Colab script to carry out the calculation.

### AlphaFold model preparation

We used the *Phenix* tool *process_predicted_model*^*14*^ to trim residues that had lower than moderate confidence (pLDDT < 70), to convert pLDDT values to estimated atomic displacement parameters, and to automatically split predicted models into domains.

### Molecular replacement

We used *Phaser*^19^ to carry out default molecular replacement (MR) analyses with X-ray data and the processed AlphaFold predictions. The number of copies of each prediction used in MR was the number of copies of the corresponding sequence in the deposited sequence file. During MR, several values of the high-resolution limit were automatically tried and the one that yielded either a result reported as convincing by the *Phaser* software^19^ or the highest value of the log-likelihood gain was used. The high-resolution limit of the data used after molecular replacement was the resolution obtained from the PDB entry.

After the first cycle, the trimmed rebuilt model from the previous cycle was used in place of a model from MR.

### Morphing predicted models onto rebuilt models

Full-length predicted models were morphed (distorted) to match the model obtained from MR using the *Phenix* tool *superpose_and_morph*^*17*^ and the keyword *direct_morph*. This tool automatically identifies parts of the predicted model that match the target model, superimposes those parts, and then smoothly deforms the model between superposed parts. This morphing procedure creates plausible models when the required distortion is small (in the range of a few Å). However, when the distortion is large, the models can be highly implausible.

### Refinement

The *Phenix refine*^*36*^ tool was used for refinement with default values for all parameters, except that checks for overlapping atoms and long bond lengths were disabled to allow refinement to proceed even if an implausible model was encountered.

### Model rebuilding

Models obtained after MR and refinement were automatically rebuilt with the *Phenix AutoBuild*^*37*^ tool, which carries out iterative crystallographic rebuilding, refinement and density modification to yield a rebuilt model and a density-modified map. The resulting density-modified map was then used in a second stage of model-rebuilding in which the model obtained from MR and refinement was rebuilt using real-space *Phenix* tools developed for models from cryo electron microscopy^17^ experiments. After carrying out these two approaches, the model that had the lowest free R value was used in subsequent steps.

### Model trimming to match maps

We trimmed the rebuilt models and docked AlphaFold predictions to provide hypotheses for further structure determination steps that only include the parts of the model that are likely to be correct. The basis for choosing which segments (sequential sets of residues) in a model to keep included map-model comparisons and, for predicted models, predicted model confidence. Map-model comparisons consisted of comparing each residue in a model to the corresponding density map (typically the density-modified map from model rebuilding) and calculating the local map-model correlation (the correlation between map values and those in a map calculated from the model). Predicted model confidence consisted of the pLDDT values from AlphaFold prediction.

The first trimming step consisted in removing segments for which the local map correlation and (for predicted models) the pLDDT values are below their corresponding cutoff values. The map correlation and the pLDDT values were smoothed over a window of typically 10 residues (controlled by the parameter *minimum_domain_length*). The cutoff values are calculated from the mean and standard deviation of the highest half of the correlation (or pLDDT) values, where the cutoff is typically the mean minus three times the standard deviation (controlled by the parameter *cc_sd_ratio*).

The second trimming step consisted in removing residues at the segment ends that have map correlation and smoothed map correlation below a higher cutoff (typically the mean minus twice the standard deviation, controlled by the parameter *cc_sd_ratio_end*). Segments that are shorter than the length of the smoothing window are then removed.

Finally, segments with a much lower mean map correlation than most segments are removed. This is achieved by calculating the average map correlation for each segment. Then, the mean and standard deviation of the top half of these average map correlations are calculated. A cutoff is then calculated as the higher of the mean correlation scaled by a constant with a typical value of 0.64 (given by the square of the parameter *reasonable_cc_ratio*), and the mean correlation minus a constant with a typical value of 0.3, given by twice the parameter *reasonable_cc_diff*). All segments with a mean map correlation below this cutoff are removed, and a new model is created from all remaining segments.

### Construction of docked AlphaFold predictions

In order to create a docked model in which each chain was identical to an AlphaFold prediction, the chains in a rebuilt model were used to guide the positioning of AlphaFold predictions. These docked predicted models are not always geometrically plausible, as the AlphaFold predictions are not necessarily the same as the corresponding structures. Rather these docked predicted models are a convenient vehicle for managing the AlphaFold predictions for a structure, and in some cases they are a reasonable representation of the structure.

### Model comparisons and map-model correlations

We compared models that were not previously superposed using least-squares superposition and calculation of rms differences in C_α_ positions with the *Phenix*^*35*^ tool *superpose_pdbs*. Models that were already placed in appropriate crystallographic positions were compared using automatic mapping of positions based on space-group symmetry using the *Phenix resolve* tool with the *compare_pdb* keyword.

Map-model correlations were calculated with the *Phenix* tool *get_cc_mtz_pdb* which maximizes local map-model correlation by (1) adjusting the radius used for masking the map around each atom and (2) modifying side-chain atomic displacement factors by adding an incremental value for each atom beyond the C_β_ atom. Correlations reported are global (using the entire map).

### Map and model display

Figures were prepared with ChimeraX^38^ version 1.2.5.

### Data Availability

Input data for deposited models were taken from the Protein Data Bank. The 215 accession codes used were: 7tzp, 7mku, 7b3n, 7tj1, 7e0m, 7tem, 7ety, 7ncy, 7qdv, 8cuk, 7o51, 7waa, 7wbk, 7tmu, 7p3a, 7wnn, 7t7j, 7ed6, 7tog, 7n0j, 7eox, 7w3s, 7Lbk, 7dtr, 7s5o, 7e85, 7tbs, 7×77, 7e1d, 7rc2, 7wja, 7tvc, 7fiu, 7nqd, 7toj, 7tfq, 7dms, 7Lw0, 7unn, 7rr3, 7etx, 7fhr, 7b1k, 7kzh, 7eus, 7twc, 7vwk, 7tok, 7f0o, 7tcr, 7v1q, 7vob, 7rm7, 7f2a, 7v6p, 7dri, 7q3a, 7Lsv, 7s0r, 7thw, 7o5y, 7mo0, 7dqx, 7vrb, 7es4, 7tt9, 7oc3, 7esi, 7u2r, 7t8o, 7pt5, 7v3b, 7dz9, 7dnt, 7r1o, 7eio, 7tcb, 7n3v, 7e3z, 7aoj, 7vnx, 7v38, 7msL, 7eyj, 7drh, 7×4e, 7e8r, 7ew8, 7t26, 7oq6, 7waw, 7vgm, 7edc, 7bLL, 7msn, 7f8e, 7b1n, 7tL5, 7t7y, 7ksp, 7Ljh, 7wdq, 7vsp, 7dn8, 7tb5, 7t8L, 7ewj, 7×8v, 7tkv, 7fit, 7sez, 7qs4, 7e6v, 7pgk, 7kdx, 7np8, 7wsj, 7mnk, 7vrc, 7mq1, 7ecd, 7dpe, 7p0h, 7exx, 7dsu, 7n29, 7ptb, 7trv, 7oom, 7ejg, 7ron, 7dxn, 7oa7, 7raw, 7w82, 7fh6, 7djj, 7vvv, 7nxg, 7mc4, 7mnp, 7vkb, 7tn3, 7mnq, 7v9h, 7s3L, 7ovp, 7f0s, 7naz, 7v9g, 7tn5, 7u02, 7smo, 7aov, 7tn6, 7v9n, 7pvh, 7zcv, 7t85, 7t7t, 7rpq, 7mkk, 7wze, 7rds, 7v6h, 7ne9, 7v9f, 7w79, 7fjg, 7fi3, 7trw, 7rw4, 7ocn, 7dn9, 7mvx, 7e4d, 7mbw, 7s5L, 7o9p, 7rLz, 7f9h, 7rpy, 7e1L, 7os0, 7f4L, 7qvb, 7e8j, 7wa4, 7v9i, 7f05, 7dpi, 7eew, 7ejw, 7b76, 7e1b, 7vmt, 7qgf, 7rqf, 7ri3, 7bgs, 7etr, 7enm, 7rb4, 7n5u, 7m4m, 7rfo, 7mqq, 7r22, 6zpp, 7mq3, 7nbv, 7t24, 7txc, 7v63, 7wwn. All models are downloadable from the PDB with links such as: https://files.rcsb.org/download/7tzp.pdb or (for larger models that are not available in this format) https://files.rcsb.org/download/7tzp.cif. We used the Phenix tool *fetch_pdb* to download models and crystallographic data for each structure.

Predicted models, rebuilt models, and density-modified map coefficients are available at: https://phenix-online.org/phenix_data, along with a spreadsheet that contains all the raw data and analyses described here. The directory terwilliger/alphafold_crystallography_2022/ contains a README file describing the contents of the site, the spreadsheet, and a data/ directory with one compressed archive for each structure containing models and crystallographic data files.

This directory also contains a compressed archive (*alphafold_crystallography*.*tgz*) containing all the data and all the scripts used to create the spreadsheet.

### Code Availability

All code for the Phenix version of the AlphaFold2 Colab is freely available on GitHub at https://github.com/phenix-project/Colabs. All code for Phenix is available at phenix-online.org.

### Additional Information

Correspondence and requests for materials should be addressed to

## Supplementary Figure

**Supplementary Figure 1.**
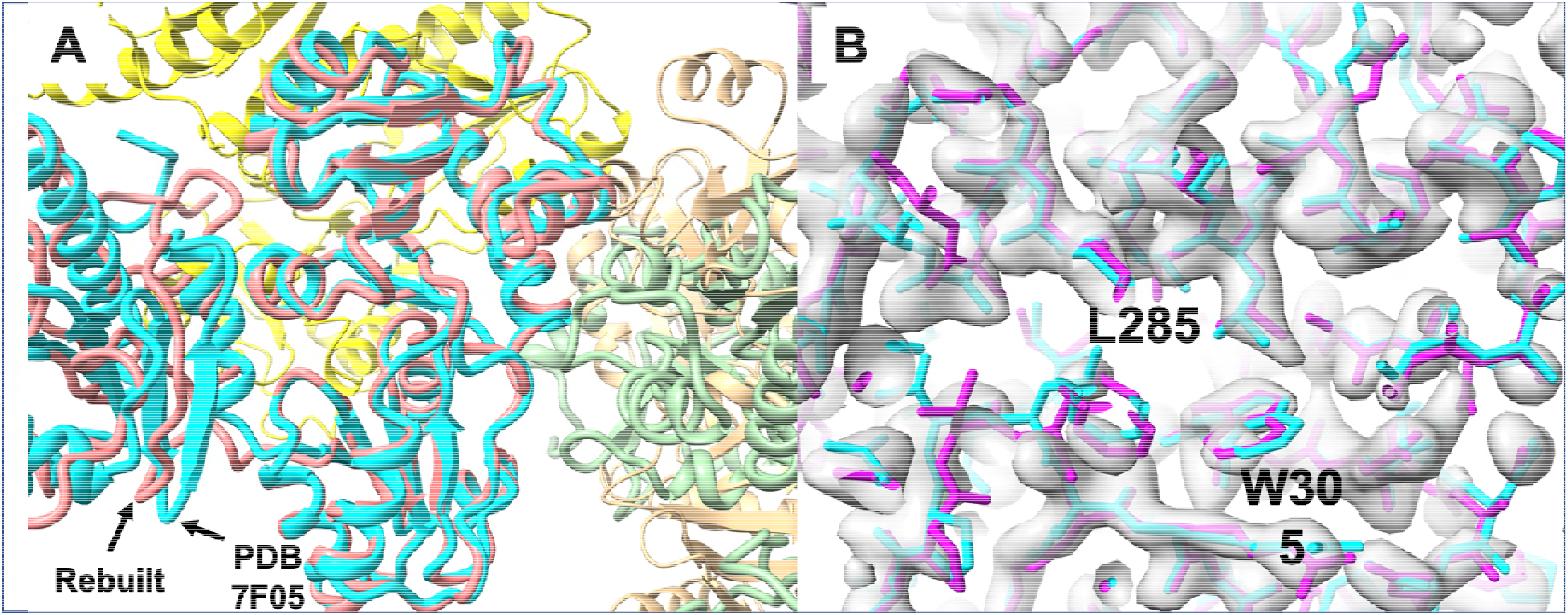
A. Rebuilt model for PDB entry 7f05 (brown, chain B, green, chain C) and deposited model (light blue, chain A, yellow, chain B, light brown, chain D, chains superposed with space group symmetry on rebuilt model). B. Density-modified electron density density map after rebuilding, rebuilt model (magenta) and deposited model (light blue) shown in the region of chain B of rebuilt model (region of chain A of deposited model).

